# Vital role for *Plasmodium berghei* Kinesin8B in axoneme assembly during male gamete formation and mosquito transmission

**DOI:** 10.1101/664375

**Authors:** D. Depoix, S.R. Marques, D.J.P. Ferguson, S. Chaouch, T. Duguet, R.E. Sinden, P. Grellier, L. Kohl

**Affiliations:** UMR 7245 CNRS Molécules de Communication et Adaptation des Micro-organismes, Muséum National d’Histoire Naturelle, Sorbonne Universités, CP52, 61 rue Buffon, 75231 Paris Cedex 05, France; Department of Life Sciences, Imperial College of London, London, United Kingdom; Nuffield Department of Clinical Laboratory Science, University of Oxford, Oxford, United Kingdom; Institute of Parasitology, Macdonald Campus, McGill University 21, 111 Lakeshore Road H9X3V9, Sainte-Anne-de-Bellevue QC, Canada

**Author notes:** Corresponding author: Delphine Depoix.

## Abstract

**Summary:** Sexual development is an essential phase in the *Plasmodium* life cycle, where male gametogenesis is an unusual and extraordinarily rapid process. It produces 8 haploid motile microgametes, from a microgametocyte within 15 minutes. Its unique achievement lies in linking the assembly of 8 axonemes in the cytoplasm to the three rounds of intranuclear genome replication, forming motile microgametes, which are expelled in a process called exflagellation. Surprisingly little is known about the actors involved in these processes. We are interested in kinesins, molecular motors that could play potential roles in male gametogenesis. We have undertaken a functional characterization in *Plasmodium berghei* of kinesin-8B (PbKIN8B) expressed specifically in male gametocytes and gametes. By generating *Pbkin8B-gfp* parasites, we show that PbKIN8B is specifically expressed during male gametogenesis and is associated with the axoneme. We created a *ΔPbkin8B* knockout cell line and analysed the consequences of the absence of PbKIN8B on male gametogenesis. We show that the ability to produce sexually differentiated gametocytes is not affected in *ΔPbkin8B* parasites and that the 3 rounds of genome replication occur normally. Nevertheless, the development to free motile microgametes is halted and the life cycle is interrupted *in vivo*. Ultrastructural analysis revealed that intranuclear mitoses is unaffected whereas cytoplasmic microtubules, although assembled in doublets and elongated, fail to assemble in the normal axonemal “9+2” structure and become motile. Absence of a functional axoneme prevented microgamete assembly and release from the microgametocyte, severely reducing infection of the mosquito vector. This is the first functional study of a kinesin involved in male gametogenesis. These results reveal a previously unknown role for PbKIN8B in male gametogenesis, providing new insights into *Plasmodium* flagellar formation.

## Introduction

*Plasmodium* parasites infect a large number of animal species and are responsible for malaria in humans. In 2017, an estimated 219 million cases of malaria and 435,000 malaria deaths were reported worldwide (World Health Organization, 2018). In most cases, malaria infections in humans are treated by artemisinin-based combination therapies, where 2 effective components with different modes of action and pharmacokinetic properties are combined. Emerging drug resistance makes it essential to search for new anti-malaria compounds and new targets in the parasite (Ouji, Augereau, Paloque, & Benoit-Vical, 2018) as no efficient vaccine for malaria is currently available.

During its life cycle, *Plasmodium* alternates between a vertebrate host and an insect vector, the female *Anopheles* mosquito. With an infective bite, the mosquito injects sporozoites in the vertebrate host, where they invade and multiply asexually, first in liver cells, and then in red blood cells. During the erythrocytic phase, a small fraction of parasites differentiate into sexual stages, male and female gametocytes. Circulating gametocytes are arrested at a G0-like stage until they are taken up by a mosquito. In the mosquito midgut, gametocytogenesis is activated when parasites are exposed to a drop in temperature, a pH increase and to xanthurenic acid (XA), a mosquito derived gametocyte-activating factor (Billker et al., 1998; Sinden, 1983; Sinden, Canning, & Spain, 1976). These changes trigger Ca^2+^ mobilization from internal stores which requires active cGMP-dependent protein kinase G (Bennink, Kiesow, & Pradel, 2016; Billker et al., 2004). While female gametogenesis involves limited morphological changes beyond escape from the host erythrocyte, male gametogenesis implicates rapid and spectacular changes (Sinden et al., 1976). In the first few minutes after activation, male gametocytes egress from the red blood cells and three rounds of rapid intranuclear DNA replication take place, resulting in an uninucleate cell with a nuclear DNA content multiplied by 8. This is accompanied by, and physically linked to, the simultaneous formation of 8 axonemes in the cytoplasm. *Plasmodium* axonemes display a classical “9+2” structure: 9 doublets of microtubules arranged in a circular pattern surrounding a central pair of singlet microtubules (reviewed in (Ishikawa, 2017)). Their mode of assembly is unusual: *Plasmodium* axonemes are assembled in the cytoplasm of the microgametocyte independent of intraflagellar transport, a bidirectional transport machinery essential for the construction of flagella in most other eukaryotes (Prevo, Scholey, & Peterman, 2017). The components of the axoneme, such as tubulin, are ubiquitous in the cytoplasm and are assembled upon activation (Kooij et al., 2005). The nuclear envelope does not break down during the 3 mitotic replications. The axonemes are however linked to the 8 replicated haploid genomes in the nucleus through spindle poles situated in nuclear pores, thus forming 8 flagellated motile microgametes, which, following violent axonemal ‘swimming’ are expelled from the microgametocyte in an exceptionally fast process called “exflagellation” (Sinden, 1983; Sinden et al., 1976). When a motile gamete encounters a female gamete, fertilization takes place resulting in a zygote. The axoneme remains intact for approximately 5 mins after fertilization, before being depolymerized in the zygote. The zygote develops into a motile and invasive ookinete, which after attachment and migration through the midgut epithelium, transforms into an oocyst. After approximately two weeks, the oocyst gives rise to sporozoites which migrate to the salivary glands and continue the parasite life cycle when injected into a new host (Bennink et al., 2016).

The sexual phase is essential to the parasite life cycle and considerable progress has been made in recent years in elucidating the molecular mechanisms governing this important differentiation process. In particular, a number of critical transcription factors and epigenetic regulators have emerged as crucial elements in the regulation of *Plasmodium* sexual commitment, allowing a better understanding of the events occurring prior to and during commitment to sexual development (Filarsky et al., 2018; Kafsack et al., 2014; Kent et al., 2018; Sinha et al., 2014). During the last years, several proteomic studies identified proteins present during the sexual stages and recently the importance of phosphoregulation was highlighted during gametogenesis (Garcia et al., 2018; Khan et al., 2005; Talman et al., 2014, Invergo et al., 2017). However, surprisingly little is known about the dramatic processes and the actors leading to free motile male gametes e.g. the molecular foundation of microgametes and how they are expelled from the microgametocyte. These dynamic processes will certainly require motor proteins, such as kinesins and dyneins, to transport cargoes, assemble the structure and generate the force for motility. Most dyneins involved in motility, are part of the complex organisation of outer and inner dynein arms present on each doublet microtubule (Oda, Abe, Yanagisawa, & Kikkawa, 2016; Roberts, Kon, Knight, Sutoh, & Burgess, 2013). In *Plasmodium*, inner dynein arms are seen less often than outer dynein arms and appear thinner in electron microscopy studies (Talman et al., 2014). This could be correlated to the absence of several inner dynein arm coding genes in the *Plasmodium* genome (Wickstead & Gull, 2007).

Only 10 kinesin encoding genes are found in the *P. berghei* genome, fewer than in other eukaryotes (Wickstead, Gull, & Richards, 2010). Among the kinesins identified by proteomic studies of sexual stages, only three are present in male gametocytes and gametes (Garcia et al., 2018; Khan et al., 2005; Talman et al., 2014). According to the comprehensive study of kinesins across eukaryotes by Wickstead et al. (2010), they belong to the families kinesin-8 (subfamily 8B), −13 and −15. Those kinesins are among the microtubule motors phosphoregulated during *P. berghei* gametocyte activation (Invergo et al., 2017). We decided to study the roles of kinesins in microgametogenesis and in particular the role(s) of kinesin-8B, the only male-specific kinesin. It will be referred to as PbKIN8B in the rest of the manuscript.

While most kinesins display predominantly one activity, either transport along microtubules or depolymerization, kinesin-8 family members are reportedly multitalented. They can walk on microtubules, but also regulate microtubule length, during mitosis (Gergely, Crapo, Hough, McIntosh, & Betterton, 2016; Grissom et al., 2009; Gupta, Carvalho, Roof, & Pellman, 2006; McHugh, Gluszek, & Welburn, 2018; Savoian & Glover, 2010), and/or ciliary and flagellar assembly (Hu, Liang, Meng, Wang, & Pan, 2015; Niwa et al., 2012; Wang et al., 2016). In *Plasmodium*, PbKIN8B could therefore be involved in several steps of microgametogenesis, such as DNA replication, axoneme assembly and function, as well as male gamete exflagellation.

Using the rodent model *P. berghei*, we show that PbKIN8B is associated with the axoneme of the microgamete. Its absence causes striking defects in axonemal structure and leads to failure to infect the mosquito and thus complete the parasite lifecycle.

## Results

### Kinesins in Plasmodium

To study kinesin motor proteins and to determine their potential role in *Plasmodium* male gametogenesis, we first compiled the known data on the proteins and their encoding genes, using the nomenclature of Wickstead et al. (2010). Nine kinesins were identified in *P. falciparum*, but the genome of several other *Plasmodium* species (*P. berghei, P. chabaudi, P. yoelii, P. knowlesi*) encodes an additional protein, kinesin-4 (PBANKA_1208200) (Table 1). *P. berghei* possesses orthologues for the kinesins identified in *P. falciparum*, either belonging to known kinesin families or currently unclassified (Table 1). Recent RNAseq studies demonstrated expression of *P. berghei* kinesins in sexual stages (male and female) (Otto et al., 2014; Yeoh, Goodman, Mollard, McFadden, & Ralph, 2017). Three kinesins were found expressed in male gametocytes and gametes by proteomic analysis (Khan et al., 2005; Talman et al., 2014): PbKIN8B, KIN13 and KIN15. As KIN13 and KIN15 are essential in the erythrocytic stage, we focussed on PbKIN8B. In *P. berghei*, PbKIN8B is composed of 1460 aa with a predicted molecular weight of 168,81 kDa and encoded by the PB_0202700 gene (4819 nt). PbKIN8B encompasses a classical motor domain positioned centrally [aa 779-1118] with 8 predicted ATP binding sites [aa 787, 872, 875, 877, 878, 879, 880, 1018] and 3 predicted microtubule interaction sites [aa 1071, 1074, 1077]. With the exception of the kinesin motor domain and of coiled-coil motives in the C-terminal region, no additional protein signatures or nuclear localisation signals (NLS) were detected.

**Table 1.**
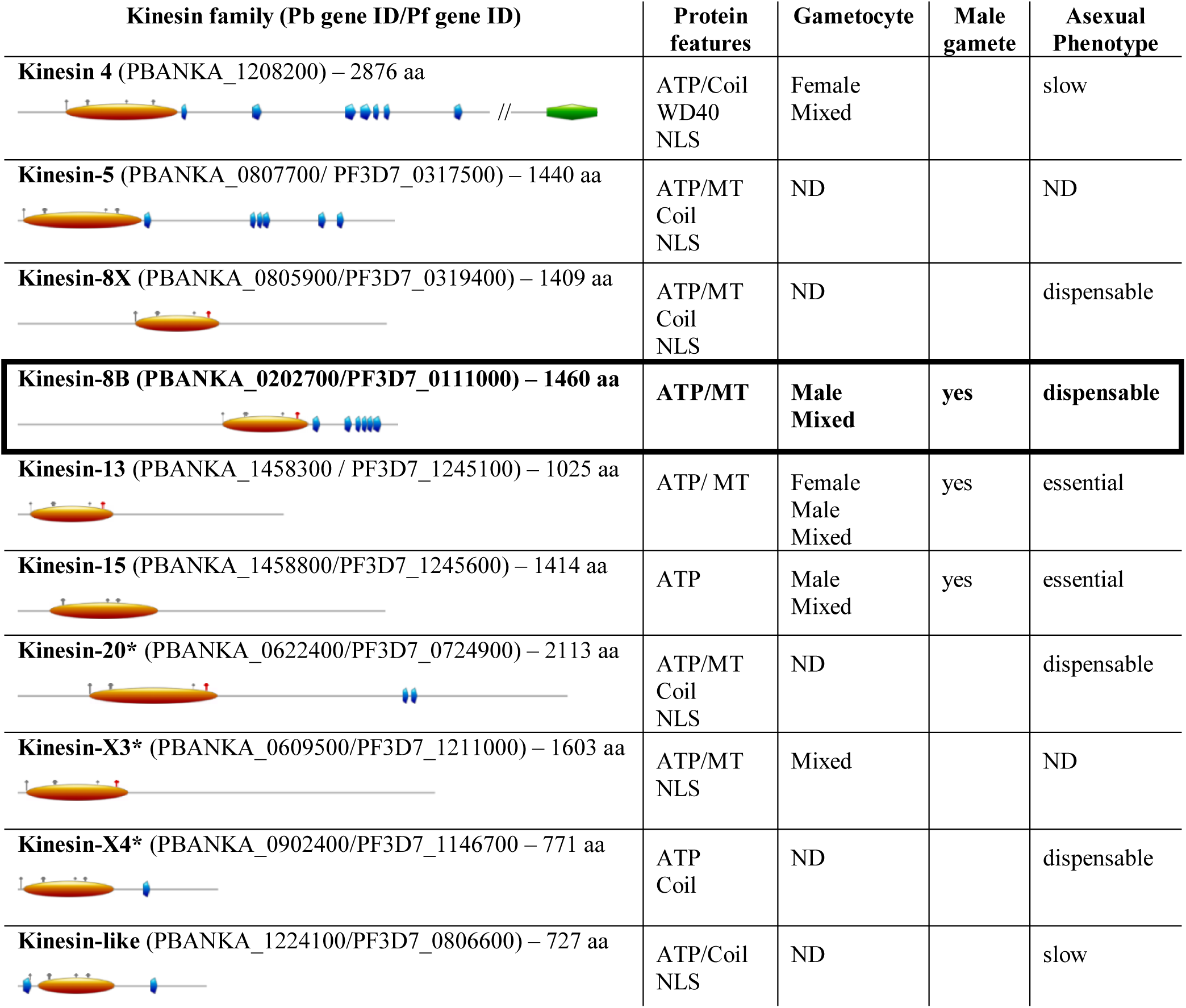
Main characteristics of *Plasmodium berghei* kinesins: protein features, proteomic data in sexual stages, phenotypic analysis in erythrocytic stages. Gene identifiers are shown for P. berghei ANKA and P. falciparum 3D7 (Plasmodb.org). Seven proteins have been associated to known kinesin families (nomenclature from (Wickstead, Gull, et al., 2010), three proteins could not be classified. Protein features were determined using PROSITE (Sigrist et al., 2013) and cNLS mapper (Kosugi, Hasebe, Tomita, & Yanagawa, 2009). In addition to the conserved kinesin motor domain with ATP binding sites and microtubule interaction (MT) sites, most proteins display coiled-coil motifs, which could be involved in protein oligomerisation. Kinesin-4 presents a WD40 motif, often implicated in protein-protein interactions. Proteomic data of purified male and female gametocytes (Khan et al., 2005), male gametes (Talman et al., 2014), and phenotypic data of asexual stages (Bushell et al., 2017) are presented when available. The asterisk indicates a difference in family affiliation between Plasmodb.org and Wickstead et al (2010) (Wickstead, Gull, et al., 2010). The characteristics of PbKIN8B are highlighted in bold. The motor domain is represented by an orange oval, ATP binding sites are shown in blue and microtubule interacting sites in red. Protein features are depicted by coloured polygons (coil-coil region in blue and WD40 region in green).

| Kinesin family (Pb gene ID/Pf gene ID) | Protein features | Gametocyte | Male gamete | Asexual Phenotype |
| --- | --- | --- | --- | --- |
| **Kinesin 4** (PBANKA_1208200) – 2876 aa | ATP/Coil<br>WD40<br>NLS | Female<br>Mixed |  | slow |
| **Kinesin-5** (PBANKA_0807700/ PF3D7_0317500) – 1440 aa | ATP/MT<br>Coil<br>NLS | ND |  | ND |
| **Kinesin-8X** (PBANKA_0805900/PF3D7_0319400) – 1409 aa | ATP/MT<br>Coil<br>NLS | ND |  | dispensable |
| **Kinesin-8B** (PBANKA_0202700/PF3D7_0111000) – 1460 aa | **ATP/MT** | **Male**<br>**Mixed** | **yes** | **dispensable** |
| **Kinesin-13** (PBANKA_1458300 / PF3D7_1245100) – 1025 aa | ATP/ MT | Female<br>Male<br>Mixed | yes | essential |
| **Kinesin-15** (PBANKA_1458800/PF3D7_1245600) – 1414 aa | ATP | Male<br>Mixed | yes | essential |
| **Kinesin-20\*** (PBANKA_0622400/PF3D7_0724900) – 2113 aa | ATP/MT<br>Coil<br>NLS | ND |  | dispensable |
| **Kinesin-X3\*** (PBANKA_0609500/PF3D7_1211000) – 1603 aa | ATP/MT<br>NLS | Mixed |  | ND |
| **Kinesin-X4\*** (PBANKA_0902400/PF3D7_1146700 – 771 aa | ATP<br>Coil | ND |  | dispensable |
| **Kinesin-like** (PBANKA_1224100/PF3D7_0806600) – 727 aa | ATP/Coil<br>NLS | ND |  | slow |

### PbKIN8B is specific to male gametocytes and gametes and is associated with the axoneme

During *Plasmodium* gametogenesis, mitosis and axoneme assembly are interconnected and happen at the same time. We wanted to know whether PbKIN8B could be involved in these processes. PbKIN8B localization was determined in the rodent model *Plasmodium berghei* by tagging the *Pbkin8B* with *gfp* at the C-terminal end. After transfection, a single crossover recombination at the endogenous PBANKA_0202700 locus resulted in a cell line expressing a *Pbkin8B-gfp* fusion protein. Two clones resulting from two independent transfections (*Pbkin8B-gfp*-cl4 and *Pbkin8B-gfp*-cl6) were isolated and the correct integration of the plasmid in the genome was confirmed by PCR (Fig S1 A, B).

The expression of the hybrid protein did not impact on the progression of the parasite asexual erythrocytic cycle. *Pbkin8B-gfp* parasites produced gametocytes in similar quantities to wt parasites with a similar sex ratio (Fig S1C, D). The parasites were transmitted to mosquitoes and naïve mice, similar to wt (see below). The *Pbkin8B-gfp* cell lines express a protein of approximately 180 kDa, recognized by an anti-GFP antibody, consistent with the calculated molecular weight of the tagged protein (*i.e* 187 kDa) (Fig 1A). The hybrid protein is expressed only in the sexual stage and is not detected in asexual parasites or in wt (Fig 1A).

**Fig 1.**
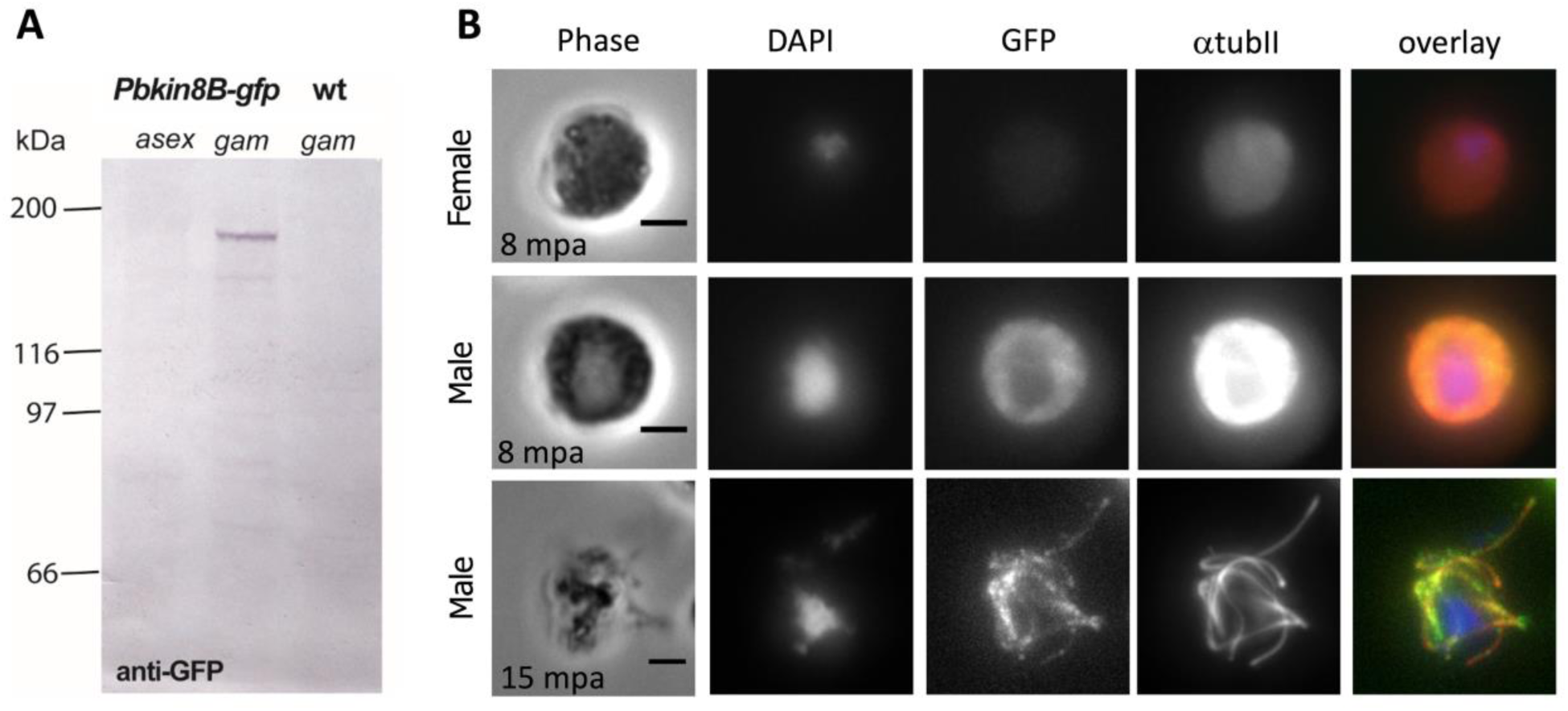
*Pbkin8B-gfp* is specifically expressed in male gametocytes and localizes to the axoneme. (A) Western blot of Pbkin8B-gfp erythrocytic stages showing expression of a 190 kDa protein only in gametocytes (Pbkin8B-gfp gam). No signal was detected in Pbkin8B-gfp asexual stages (Pbkin8B-gfp asex) nor in wt gametocytes (wt gam). Similar results were obtained with both clones. (B) Immunofluorescence assay of Pbkin8B-gfp male and female gametocytes at 8 minutes post activation (mpa) and exflagellating male gametes at 15 mpa. DAPI staining of DNA is seen in blue, anti-α tubulin II in red and anti-gfp in green. Immunofluorescence images correspond to the maximum intensity projection of the z-series. Gfp images were treated post capture using ImageJ for better visualisation (increase by 2 times for gfp gametocytes and 10 times for exflagellating microgametes). The colocalisation of α tubulin II and gfp evidence male specific expression of Pbkin8B-gfp. Scale bar: 2 µm.

We then studied the cellular localization of PbKIN8B, by following the expression of *Pbkin8B-gfp* by IFA at different stages of the life cycle (Fig 1B). Male and female gametocytes can be easily distinguished during the course of gametogenesis because amount of tubulin (seen by the marker α–tubulin II (Kooij et al., 2005)) and DAPI intensity vary: they increase in male gametocytes, while remaining almost constant in female gametocytes. *PbKIN8B-gfp* could not be detected in asexual parasites, nor in female gametocytes.

In unactivated male gametocytes, the protein shows a uniform cytosolic distribution, with the exclusion of the nuclear region. Upon activation by XA, *in vitro*, the localization of the *Pbkin8B-gfp* signal changes and is seen as punctiform lines with spatial and temporal dynamics similar to the axonemal marker α–tubulin II (Fig 1B).

After exflagellation, the *Pbkin8B-gfp* signal is detected in the male gametes, but it can not be detected after fertilization in ookinetes (data not shown). This could be due to a diffuse localization of the protein in the ookinete and the oocyst, hindering its detection under the conditions used for IFA or an absence of PbKIN8B at these stages.

### PbKIN8B plays a crucial role in exflagellation and fertilization *in vitro*

Several potential roles for PbKIN8B in male gametogenesis could be envisaged: it could be involved in mitosis, flagellum assembly/functioning and/or exflagellation. We generated *P. berghei* PbKIN8B knock-out parasites (named *ΔPbkin8B*) by double homologous recombination using the PlasmoGem construct PbGEM-267699 replacing the PbKIN8B encoding gene by a human *dhfr/ts* cassette (Fig S2 A-C). After selection by pyrimethanime and cloning by limiting dilution, two *ΔPbkin8B* clones, resulting from two independent transfections (*ΔPbkin8B*-cl3600 and *ΔPbkin8B*-cl3716), were obtained. Both cell lines show gametocyte numbers and sex ratio similar to wt parasites (Fig S2 D, E).

The *ΔPbkin8B* mutant parasites are able to form asexual and sexual blood stages, therefore mitosis in asexual blood stages is normal. The defects in the *ΔPbkin8B* mutant parasites are therefore restricted to microgametogenesis.

In blood containing wt gametocytes, induction by XA leads to the release of male gametes from microgametocytes within 15 min post activation (mpa), as well as adherence to surrounding red blood cells forming “exflagellation centres”. In *ΔPbkin8B* parasites no exflagellation centres were observed in 10 experiments (5 per clone), even after a prolonged induction period (up to 30 mpa). This contrasts strikingly with exflagellation in wt parasites (33,46 ± 17,87 exflagellation centres per 100 male gametocytes, at 15 mpa) (Fig 2A). No free male gametes were observed in the mutant lines. Recognising that rare gametes could have been missed in the mutant even with careful observation, and that an isolated male gamete could be sufficient to fertilise a female gamete, we performed an *in vitro* ookinete conversion assay to quantify the proportion of activated female gametes that converted into ookinetes. In wt parasites, the mean ookinete conversion rate was 63.5 ± 3.6, whereas it was only 1.2 ± 1.4 and 0.2± 0.2 for *ΔPbkin8B* clones 3600 and 3716 (Fig 2B), indicating that a few viable male gametes were produced in the mutant that were able to fertilize the female gametes.

**Fig 2.**
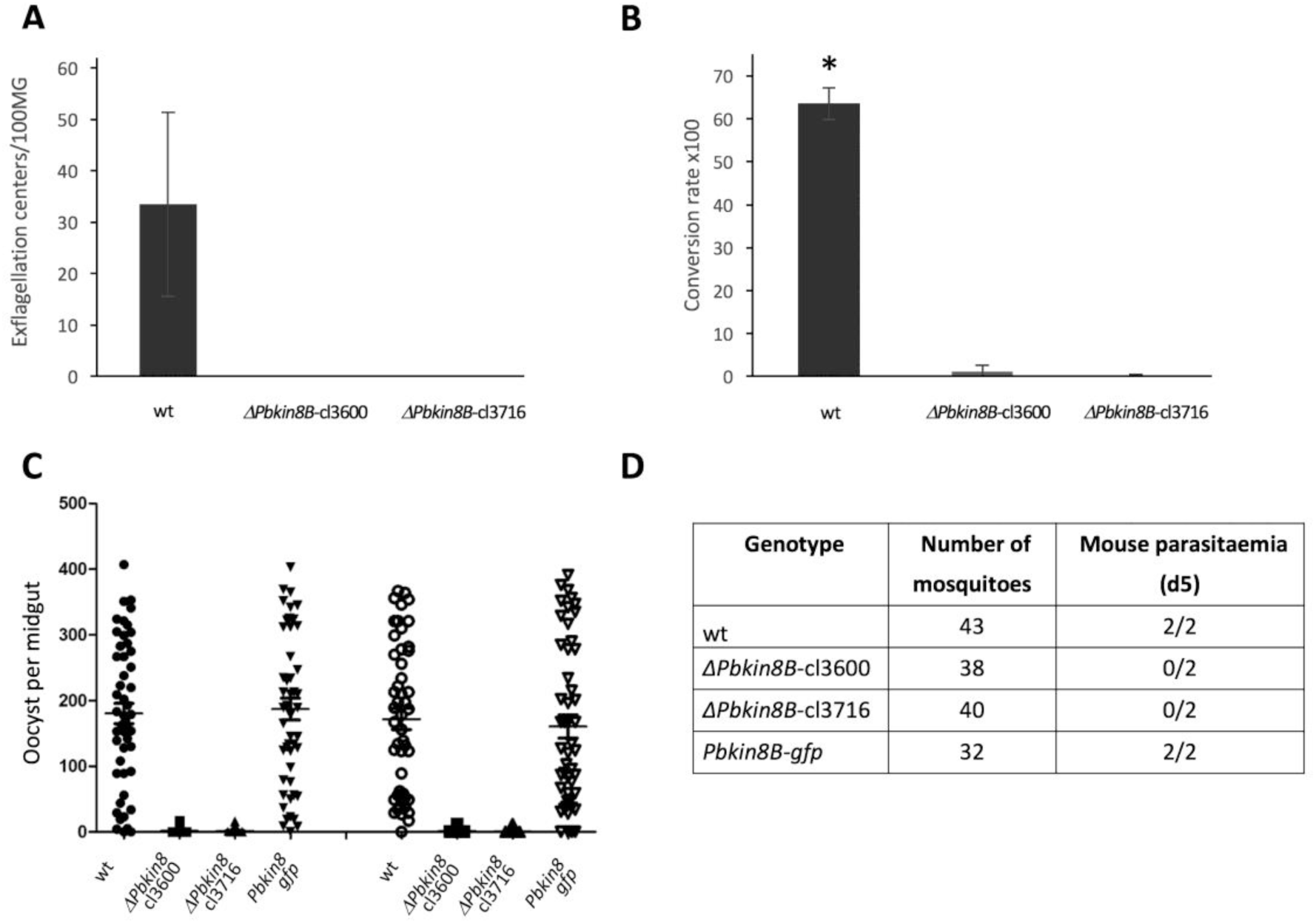
*ΔPbkin8B* parasites are unable to complete the parasite life cycle. (A, B) In vitro analysis of the ΔPbkin8B mutant. (A) Comparison of exflagellation rates between ΔPbkin8B (clones 3600 and 3716) and wt cells at 15 mpa. Numbers of exflagellation centres are expressed relatively to 100 microgametocytes. (B) Comparison of ookinete conversion rates between ΔPbkin8B (clones 3600 and 3716) and wt. For each replicate (n=5), the conversion rate was calculated as the percentage of macrogametes that developed into ookinetes. Values represent an average of more than 800 cells. SD are reported as bars on the figures. Asterisk (Ȫ) indicate statistically significant differences in Student’s T-test with p-values lower than 0,01. (C, D) In vivo analysis of the ΔPbkin8B mutant (2 clones). (C) Comparison of mosquito infection rates between ΔPbkin8B (clones 3600 and 3716), Pbkin8B-gfp and wt parasites. At day 8 and day 12 post feeding, infected A. stephensi mosquitoes were dissected and oocysts, were counted in each midgut. Detailed results are presented in Supplemental table 2. (D) Analysis of transmission of ΔPbkin8B, Pbkin8B-gfp and wt parasites from mosquitoes to naïve mice. A. stephensi mosquitoes infected respectively with ΔPbkin8B, Pbkin8B-gfp and wt parasites, were allowed to bite naïve mice. Infection of mice was monitored at day 5 and day 12 on Giemsa stained blood smears.

### PbKIN8B is essential for the completion of the life cycle *in vivo*

*In vitro* experiments do not necessarily reflect the complexity of the parasite life cycle, therefore we analysed the life cycle *in vivo* by transmitting wt and mutant parasites to mosquitoes and naïve mice. We determined the capacity for infection of *ΔPbkin8B* (2 independent clones), wt and *Pbkin8B-gfp* parasites. *A. stephensi* mosquitoes were allowed to feed on mice infected with the different parasite lines. Eight and twelve days post feeding, the midgut of the mosquitoes was dissected and the number of oocysts per midgut was determined (2 independent experiments). The two *ΔPbkin8B* mutant cell lines produced very few oocysts (mean range of 1.68/1.36 and 1.4/0.76 respectively) in comparison to wt and *Pbkin8B-gfp* parasites, where numerous oocysts were formed (mean values the 2 replicates of 180/171 and 187/160 oocysts per midgut respectively) (Fig 2C, Table S2). No sporozoites could be detected in the *ΔPbkin8B* oocysts under microscopy.

**Table 2.**
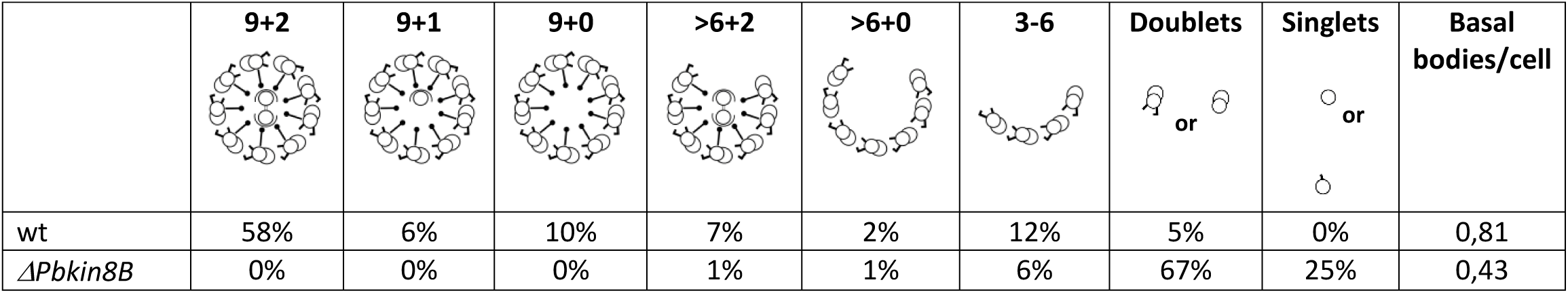
Quantification of microtubule structures and basal bodies in *ΔPbkin8B* and wt gametocytes. Arrangement of doublet microtubules in circles, with or without a central pair of microtubules, semi-circles or isolated doublet, respectively singlet microtubules, were counted in 47 ΔPbkin8B and 18 wt gametocytes and the results are presented as percentage of each type. The basal body counts are presented as relative number of basal bodies per cell observed in random thin sections through microgametocytes based on examination of 27 wt and 23 ΔPbkin8B cells.

Since as few as 10 sporozoites would be sufficient to complete the life cycle (Churcher et al., 2017), we tested the ability of the parasites to infect mice. *A. stephensi* mosquitoes, fed 21 days prior on mice infected with *ΔPbkin8B*, wt or *Pbkin8B-gfp* parasites, were allowed to bite naïve 6-to 8-week-old female Tucks Ordinary mice. Wt and *Pbkin8B-gfp* fed mosquitoes were able to transmit parasites to the naive mice, as observed on Giemsa stained blood smears at day 5 post feeding. In contrast, mosquitoes fed with *ΔPbkin8B*-cl3600 and cl3716 were unable to transmit parasites to naive mice (Fig 2D). Parasitaemia was followed up to 14 days after feeding in the mice bitten by mosquitoes with *ΔPbKIN8B* parasites, but no parasites were seen.

### Dissection of the role of PbKIN8B in male gametogenesis

The *ΔPbkin8B* mutant displays little evidence of microgametes and is unable to complete the parasite life cycle, indicating a major defect in male gametogenesis. As this process is composed of different events such as host cell egress, DNA replication and axoneme formation, and finally exflagellation, we investigated each of these steps in the *ΔPbkin8B* mutant at different times post activation.

First, we followed cell egress in *ΔPbkin8B* mutant cells and wt cells, using an anti-spectrin antibody, which recognizes the most abundant protein in the red blood cell membrane skeleton. After induction by XA, both *ΔPbkin8B* and wt gametocytes were able to escape from the host cell membrane as shown by the weaker signal with the anti-spectrin antibody (Fig S3).

Second, we looked at the 3 rounds of endomitoses that occur consecutively during microgametogenesis resulting in an increase of the DNA content. In IFA, we were not able to see a localisation of *PbKIN8B-gfp* in the nucleus. During *Plasmodium* gametogenesis, mitosis and axoneme assembly are interconnected and happen in the same time. Defects in one process could influence the other. We therefore analysed DNA content in at least 30000 gametocytes at 0, 8 and 15 mpa by XA, using flow cytometry (Fig 3A-C). We compared *ΔPbkin8B* gametocytes to wt and *Δcdpk4* gametocytes, a mutant unable to undergo DNA replication (Billker et al., 2004). Gating was established within the YOYO+ population (P1). At time 0 min, in all cell lines, most gametocytes (male and female) possess a nuclear content corresponding to the P2 population (Fig 3A). Only a few cells with higher DNA content were detected in wt and *ΔPbkin8B* cells. During the course of the induction, the P2 population decreases slightly while the P3 population (highest DNA content cells) increases, indicating that the *ΔPbkin8B* microgametocytes replicate their DNA similar to wt, both in quantity and time frame (Fig 3B, C). The proportion of cells with highest DNA content in the sample (ratio P3/P1) increases by a factor of approximately 3 in wt and 4 in *ΔPbkin8B*. As expected, no change in DNA profile is observed in *Δcdpk4* cells over the time of the experiment (Fig 3C).

**Fig 3.**
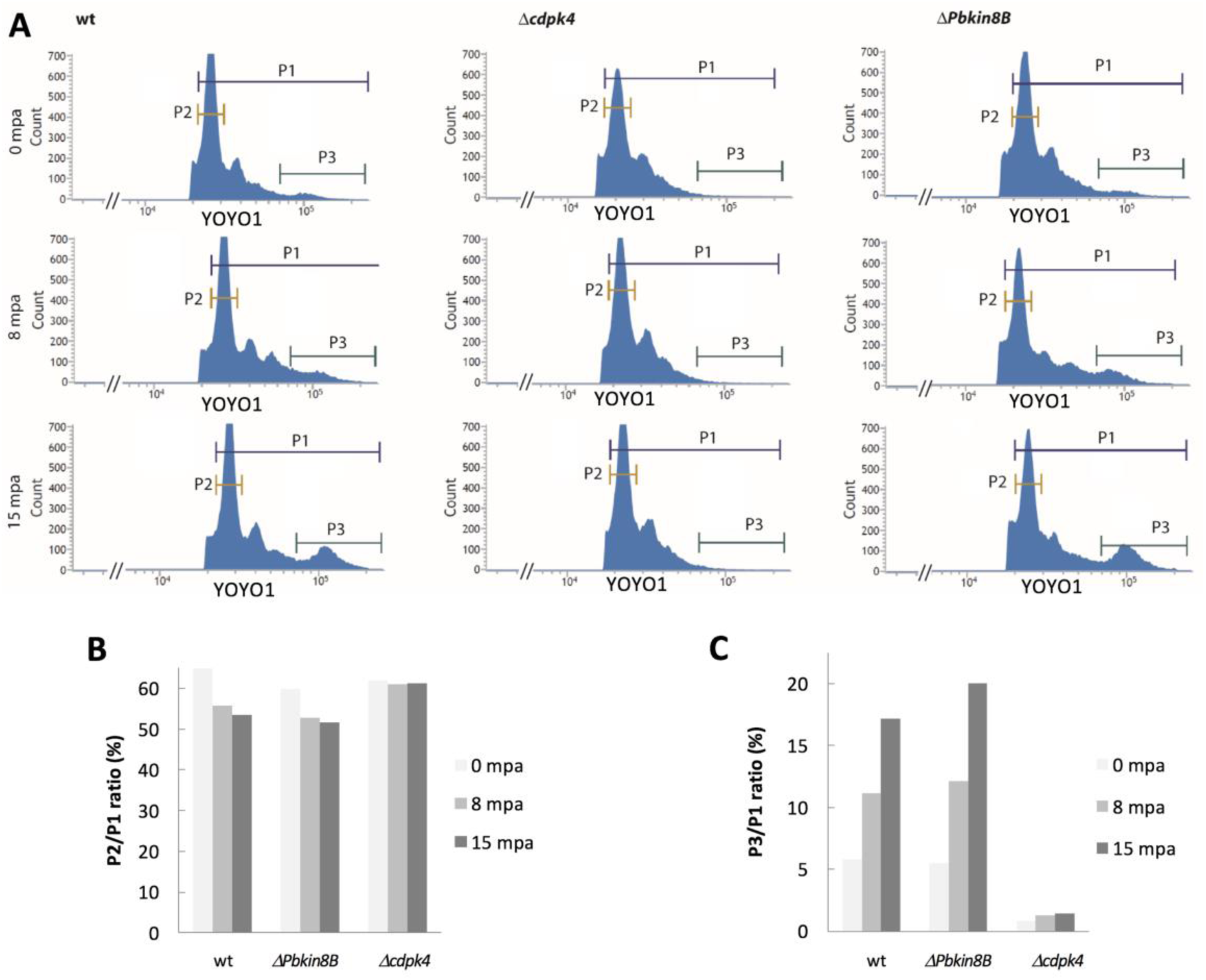
*ΔPbkin8B* cells are able to replicate their DNA during male gametogenesis. (A) Time course analysis of gametocyte DNA content by flow cytometry. Purified mixed gametocytes from ΔPbkin8B, Δcdpk4 (unable to undergo DNA replication) and wt parasites were fixed at times 0, 8 and 15 mpa and DNA was labelled with YOYO-1. Gates were established defining populations P1 (total YOYO+ population), P2 (low DNA content cells), and P3 (high DNA content cells). (B, C) Evolution over time of populations P2 and P3. Proportion of P2 cells (B), respectively P3 cells (C), relative to YOYO+ cells at times 0, 8 and 15 mpa.

Third, we analysed axoneme formation by IFA using an anti α-tubulin II antibody, combined with DAPI staining (Fig 4). From the beginning of activation to 8 mpa, *ΔPbkin8B* and wt gametocytes were indistinguishable by DAPI staining: the DNA positive area became enlarged and the intensity of the signal increased, suggesting that nuclear DNA was replicated in *ΔPbkin8B* parasites similar to wt cells, corroborating the FACS results (Fig 3). At the same time, microtubules were assembled in the cytoplasm, coiled around the enlarged nucleus, though the signal in *ΔPbkin8B* cells looked less intense than in wt cells (Fig 4). At 15 mpa, the replicated DNA separated into 3-8 ‘clumps’ in wt and mutant. At this moment, the overall cell shape differed between mutant and wt parasites. *ΔPbkin8B* microgametocytes remained rounded. Short thin protrusions, labelled with the anti α-tubulin II antibody, could be seen in some of these cells (Fig 4), but they did not separate from the microgametocyte even at 30 mpa. By contrast, at 15 and 30 mpa, exflagellating gametes could be observed in the wt.

**Fig 4.**
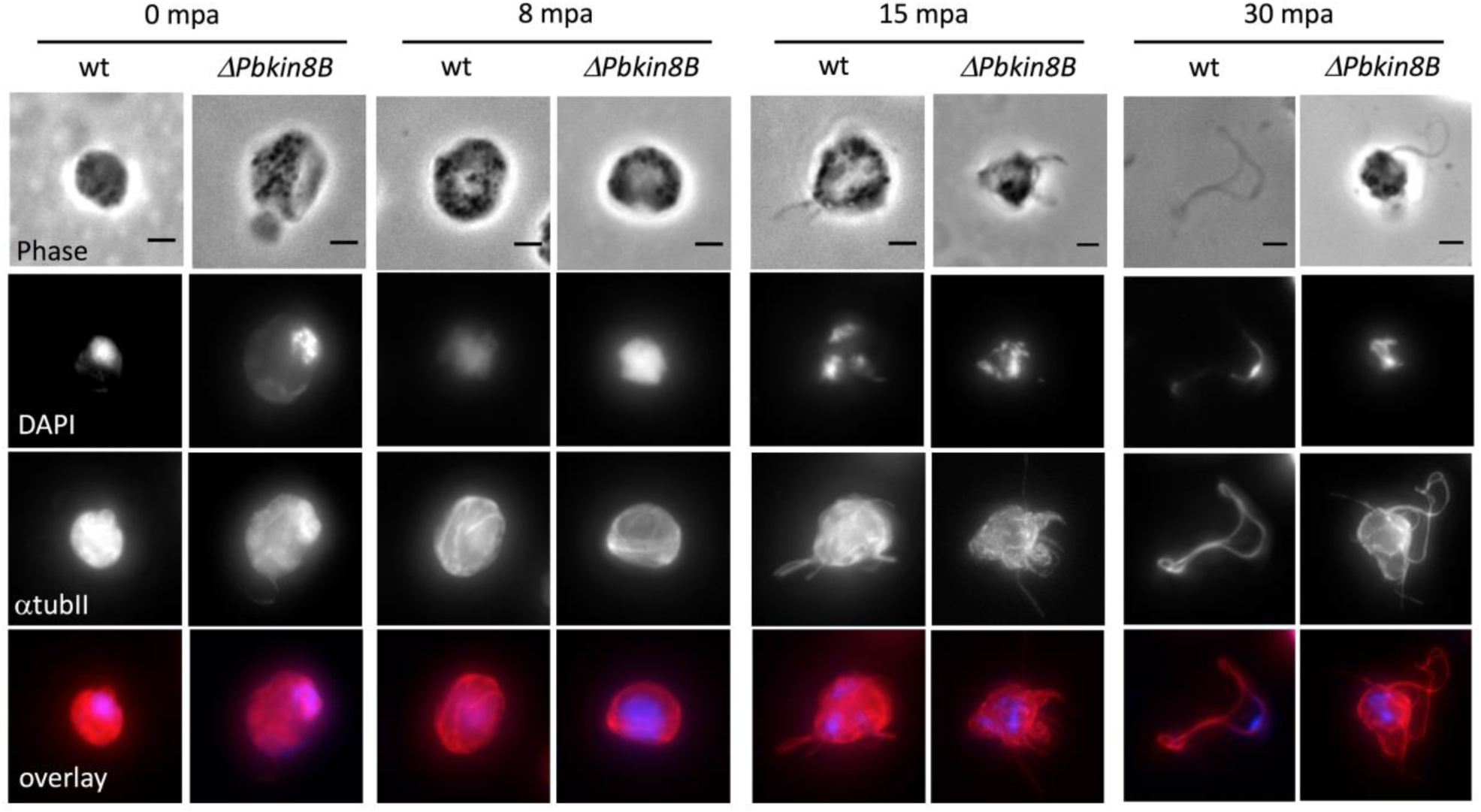
*Δ Pbkin8B* parasites are able to replicate DNA and assemble microtubules, but cannot release male gametes. Immunofluorescence assay of wt and ΔPbkin8B male gametocytes from beginning of activation to 30 mpa. DAPI staining of DNA is seen in blue and anti-α tubulin II in red. Immunofluorescence images correspond to the maximum intensity projection of the z-series. During the course of activation, wt microgametocytes increase nuclear content and formed microgametes. ΔPbKIN8B microgametocytes also replicate DNA and assemble microtubules in the cytoplasm. However, they are not able to release microgametes. Scale bar: 2 µm.

PbKIN8B is therefore involved most likely in axoneme assembly/stabilisation and/or exflagellation.

### PbKIN8B is essential for axoneme assembly in *Plasmodium*

To better understand the phenotype observed by light microscopy, we examined the ultrastructure of male gametocytes by transmission electron microscopy in wt and *ΔPbkin8B* cells at 15 and 30 mpa. At 15 mpa, the gametocytes from wt and *ΔPbkin8B* have egressed from the RBC and were observed at various stages of microgametogenesis (Fig 5A-J). The earlier stages of both wt (Fig 5A, G) and mutant (Fig 5B, H) are characterized by a large spherical central nucleus with dispersed chromatin. When the cytoplasm was examined, the wt cells presented a number of developing axonemes with classical 9 doublet microtubules around 2 central microtubules, 9+2 (Fig 5A, C, D). The microtubules of the axonemes develop from an electron dense basal body and elongate round the periphery of the microgametocyte (Fig 5A, G, I). In wt, cross section showed that the majority of axonemes were normal (60%, 9+2 based on 18 randomly selected microgametocytes), but a proportion (40%) showed varying degrees of abnormality (Table 2). In contrast, in the mutant, examination of a random sample of 47 microgametocytes failed to identify any complete “9+2” axonemes (Table 2). Organized interaction between doublet microtubules was rare although numerous doublet and single microtubules were identified in the cytoplasm (Table 2). Similar to wt, the microtubules in the mutant appear to grow from electron dense structures (basal bodies) (Fig 5B, H, J). Although difficult to quantify, there appeared to be fewer basal bodies (0.81 basal bodies per section through wt microgametocytes compared to 0.43 for mutant (Table 2)) and these appear to have lost their close connection to the nuclear pole observed in the wt (90% wt to 20% mutant) (Table S3). In cross section, the microtubules appeared to be randomly distributed in the cytoplasm (Fig 5E, F) but in longitudial sections, microtubules still appeared to elongate round the periphery of the gametocyte (Fig 5B, H). It was possible to identify two single microtubules similarly arranged to the central pair of microtubules of the axonemes but little evidence of spatial organization of the doublet microtubules around them was observed (Fig 5F). As the microgametocytes develop (undergo genome replication), multiple nuclear poles were observed in both the wt and mutant (Fig 5G, H). The organization of nuclear poles was similar in both wt and *ΔPbkin8B* consisting of an electron dense cone-like structure from which spindle microtubules radiate with attached kinetochores (insets in Fig 5G, H).

**Fig 5.**
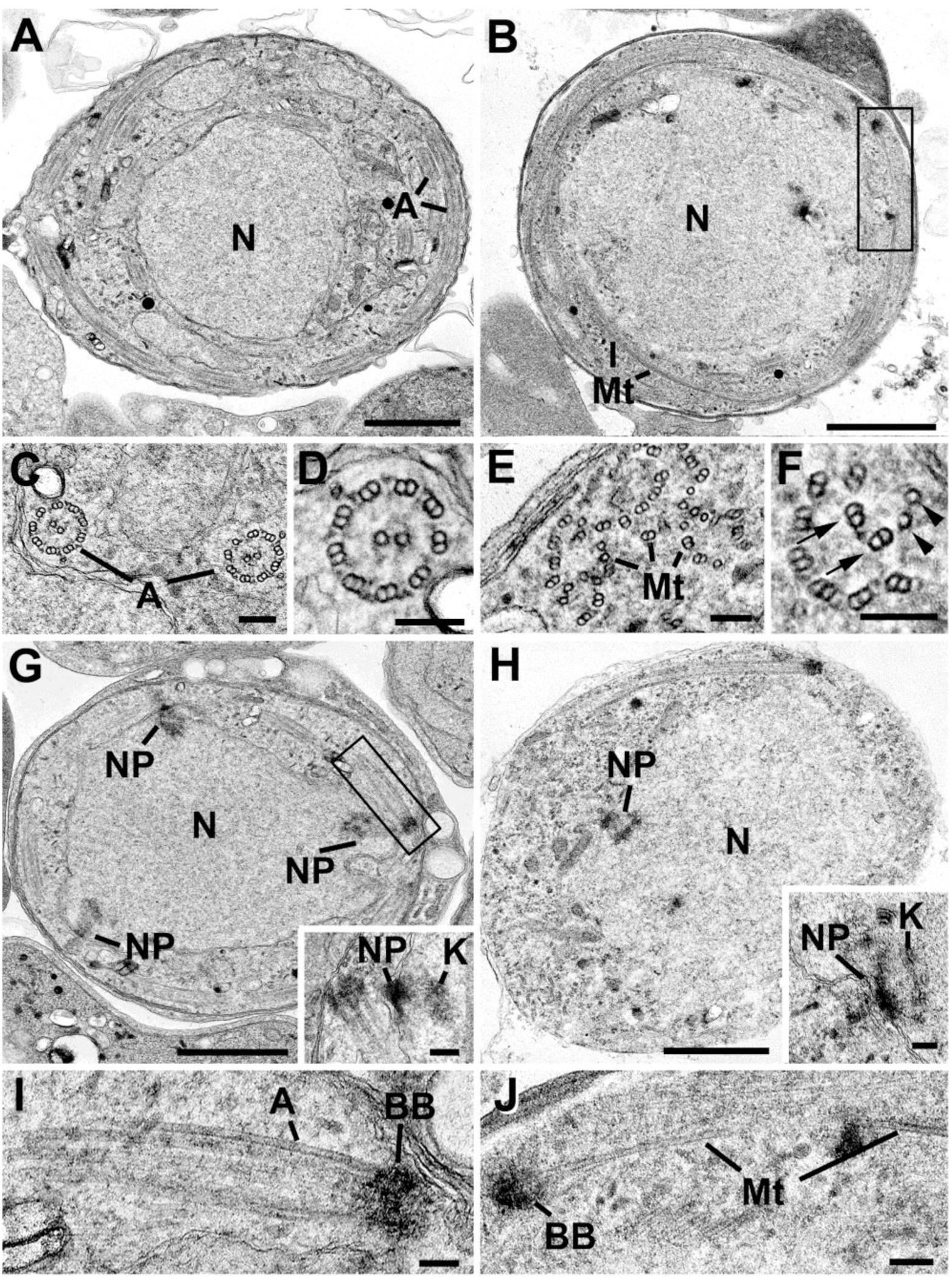
*Δ Pbkin8B* microgametocytes display a modified ultrastructure with disorganised axonemes. Electron micrographs of various developmental stages of microgametogenesis in wt and ΔPbkin8B parasites at 15 and 30 mpa. Bars represent 1µm in A, B, G, H and 100 nm in all other micrographs. A. Low power through a wt microgametocyte showing the central nucleus (N) with a number of axonemes (A) running round the peripheral cytoplasm. B. Low power through a ΔPbkin8B microgametocyte showing the large central nucleus (N) with longitudinally running microtubules (Mt). C. Detail of the cytoplasm of wt microgametocyte illustrating cross section through two axonemes (A). D. Enlargement of an axoneme showing the 9+2 arrangement of the microtubules. E. Detail of the cytoplasm of a ΔPbkin8B gametocyte showing a cross section through a number of randomly distributed microtubules seen mostly as doublets (Mt). F. Enlargement of a group of microtubules showing two single microtubules (arrowheads) with randomly distributed doublet microtubules (arrows). G. Low power of a mid-stage wt microgametocyte showing the central nucleus (N) with a number of nuclear poles (NP). Insert. Detail of a nuclear pole (NP) and radiating microtubules with attached kinetochore (K). H. Low power of a mid-stage ΔPbkin8B gametocyte showing the central nucleus (N) and associated nuclear pole (NP). Insert. Detail of a nuclear pole (NP) and radiating microtubules with attached kinetochore (K). I. Detail from the cytoplasm of a wt parasite from the enclosed area g showing a longitudinal section through the basal body (BB) and axoneme (A).J. Detail from cytoplasm of a ΔPbkin8B gametocyte from the enclosed area in b showing longitudinally running microtubule (Mt) emanating from an electron dense structure, possibly a basal body (BB).

In the later stages of microgametogenesis, in both the wt and *ΔPbkin8B* cells, there was similar chromatin condensation within the nucleus prior to potential exflagellation (Fig 6A, B). Exflagellation with a flagellum and associated nucleus protruding from the microgametocyte (Fig 6C) and free microgametes could be observed in the wt (Fig 6E, F), but no free microgametes were seen in the *ΔPbkin8B* samples although very rare examples of cytoplasmic protrusions were identified (Fig 6D), which contained disorganized doublet microtubules (Fig 6G)

**Fig 6.**
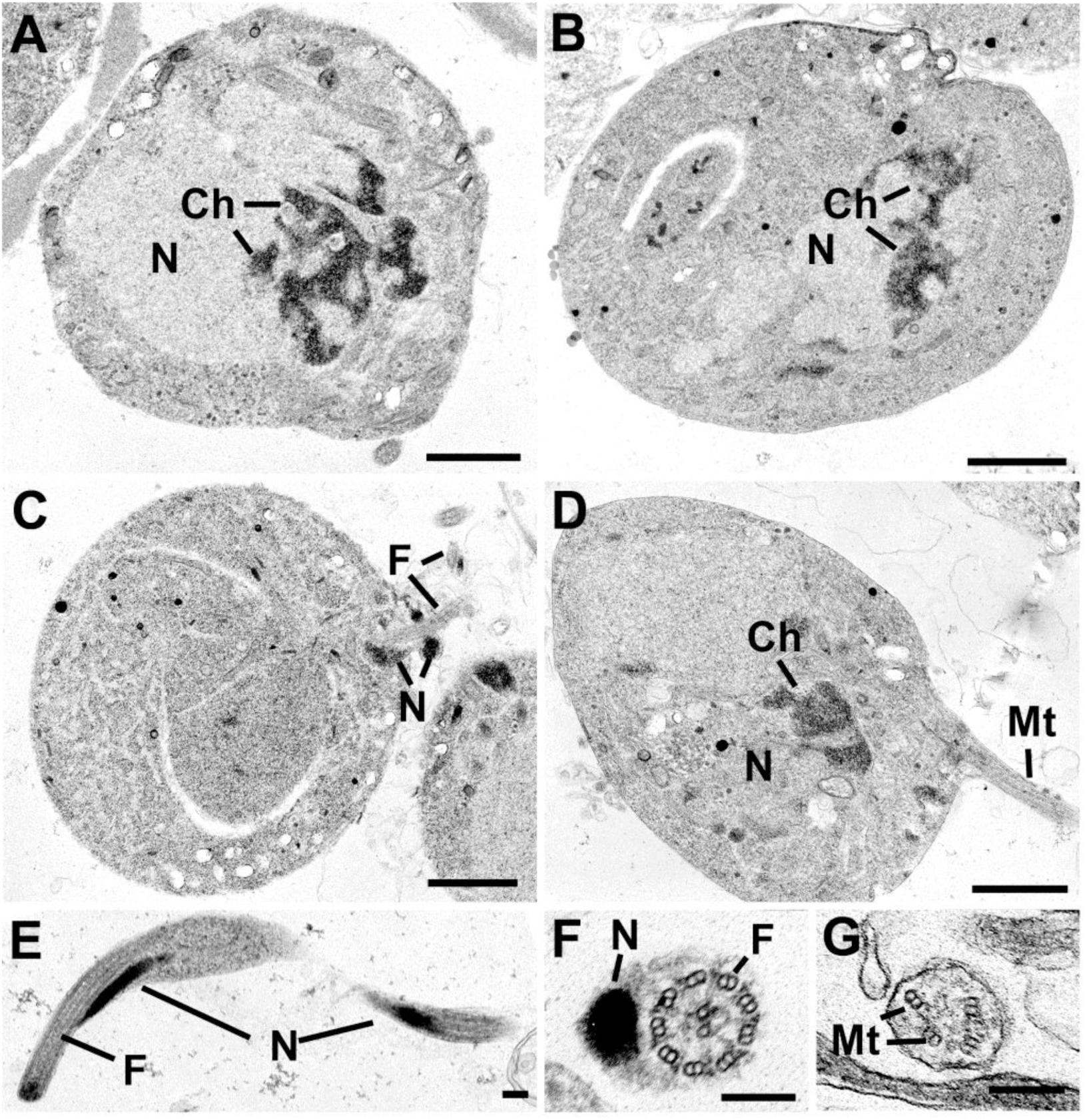
*ΔPbkin8B* microgametocytes condenses chromatin but do not release male gametes. Electron micrographs of late developmental stages of microgametogenesis in wt and ΔPbkin8B parasites at 15 and 30 mpa. Bars represent 1µm in A, B, C and D and 100 nm in all other micrographs. A. Low power of a late stage wt microgametocyte showing the nucleus (N) with peripherally condensed chomatin (Ch). B. Low power of a late ΔPbkin8B gametocyte showing the nucleus with condensed chromatin (Ch). C. Low power of a late microgametocyte undergoing exflagellation with the flagellum (F) and associated nucleus (N) protruding from the surface. D. Low power of a late ΔPbkin8B gametocyte showing a centrally located nucleus (N) with condensed chromatin (Ch) and a cytoplasmic process from the surface containing microtubules (Mt). E. Longitudinal section through a free microgamete showing the relationship between the nucleus (N) and the flagellum (F). F. Cross section through a free wt microgamete illustrating the electron dense nucleus (N) and associated flagellum (F). G. Detail of a cross section through a cytoplasmic process from a ΔPbkin8B gametocyte showing disorganized duplet microtubules (Mt).

In summary, PbKIN8B does not play a role in genome replication and intranuclear subdivision in gametogenesis, but it is essential for assembly of structured “9+2” axonemes in the cytoplasm. In the absence of viable/motile microgametes, parasites are unable to successfully complete their life cycle.

## Discussion

We hypothesized that motor proteins, such as kinesins, are essential actors in the dynamic processes of microgametogenesis in *Plasmodium*. *P. berghei* possesses members of the kinesin families 4, 5, 8, 13, 15 and 20, as well as three kinesins which could not be associated to known families (Wickstead, Gull, et al., 2010) (Table 1). There is no ortholog of kinesin-4 family member PBANKA_1208200 in *P. falciparum*, even though it is functional and important for growth at the asexual stages in *P. berghei* (Bushell et al., 2017). The restricted number of kinesin encoding genes does not necessarily mean that *Plasmodium* needs these proteins less than other organisms. Malarial kinesins could have multiple roles as it has been shown in other organisms (Dawson et al., 2007; Wickstead, Carrington, Gluenz, & Gull, 2010).

According to two transcriptomic studies, all kinesin encoding genes in *P. berghei* are expressed during the sexual stage (Otto et al., 2014; Yeoh et al., 2017). Proteomic studies identified 3 kinesins expressed during male gametogenesis (kinesin-8B, −13 and −15) (Khan et al., 2005; Talman et al., 2014), but no functional studies have been conducted until now. In other eukaryotes, orthologues for kinesin-4, −5, −8B, −13 and −15 are known to be involved in microtubule dynamics and particularly mitosis and cilia/flagella length regulation (Almeida & Maiato, 2018; Hu et al., 2015; Ma, Wang, & Yang, 2017; Muhia et al., 2016; Niwa, 2015; Niwa et al., 2012; Shrestha, Hazelbaker, Yount, & Walczak, 2018; Singh, Pandey, Al-Bassam, & Gheber, 2018; van Riel et al., 2017). Unfortunately, concerning PBANKA_0622400, PBANKA_0609500, PBANKA_0809500, PBANKA_0902400 and PBANKA_1224100, no functional data are available and the number of species possessing orthologues of these proteins is currently very limited, making it difficult to construct any hypothesis concerning their roles.

In other eukaryotes, kinesin-8 family members, which share a family-specific helical neck region (Miki et al., 2005), have been shown to be involved in both mitosis and microtubule regulation in cilia/flagella. The members of the kinesin-8A subfamily are found in different organisms and their roles in mitosis, often promoting microtubule catastrophe, have been studied extensively (Dave et al., 2018; Edzuka & Goshima, 2019; Gergely et al., 2016; Grissom et al., 2009; Messin & Millar, 2014; Savoian & Glover, 2010; Su et al., 2013). In fission yeast, additional roles involving cell polarity and control of cortical microtubule length have been shown (Meadows et al., 2018; West, Malmstrom, Troxell, & McIntosh, 2001). In contrast, members of kinesin-8B subfamily have been poorly studied. They are restricted to ciliated/flagellated organisms (Wickstead, Gull, et al., 2010), but absent in some species, like *Drosophila, Trypanosoma* or *Chlamydomonas*, inferring a particular flagellar role for these proteins. The functional information arises from the mammalian orthologue, KIF19A, localized at the ciliary tip. HsKIF19A plays a role in controlling microtubule length in cilia and its absence results in the formation of abnormally long cilia, causing hydrocephaly and female infertility in mice (Niwa et al., 2012).

*Plasmodium* possesses 2 members of the kinesin-8 family: PbKIN8B and KIN8X (PBANKA_0805900, belonging to neither the KIN8A nor the KIN8B subfamily (Wickstead, Gull, & Richards, 2010). Recently, Zeeshan et al. (2019) showed that KIN8X is localized at the mitotic spindle during cell division and that it fulfils an essential role in the parasite life cycle (Zeeshan, Shilliday, et al., 2019). Only PbKIN8B is specifically expressed in male gametocytes and gametes (Khan et al., 2005; Talman et al., 2014). We hypothesised that PbKIN8B could play multiple roles in microgametogenesis, either in mitosis and/or axoneme construction.

If PbKIN8B was involved in mitosis, a nuclear localization of the protein, as well as defects in DNA replication in its absence, might have been expected. In *Plasmodium*, as opposed to metazoan eukaryotes, the nuclear envelope never breaks down. Cytoplasmic kinesins therefore never get access to the spindle during mitosis, and nuclear *vs.* cytoplasmic location is critical. A nuclear localisation signal could not be predicted in the PbKIN8B sequence. This is in agreement with the IFA data showing that PbKIN8B is excluded from the nucleus. Moreover, in the absence of PbKIN8B, DNA replication and spindle activity occur normally in microgametes and asexual blood stage parasites. A mitotic role of PbKIN8B during other life stages where mitosis also occurs, such as sporozoite formation and/or liver-stage development, whilst not anticipated, cannot be completely excluded. Unfortunately, no data are available on the expression of PbKIN8B in these stages.

Looking at the different steps in male gametogenesis, we have shown that mutant *ΔPbkin8B* microgametocytes are able to egress from the red blood cell, a step which is independent from DNA replication and axoneme motility (Tewari, Dorin, Moon, Doerig, & Billker, 2005). The absence of PbKIN8B causes severe defects in axonemal assembly. While numerous singlet and doublet microtubules are present in the cytoplasm, they are never organised into “9+2” axonemes. *In vivo*, PbKIN8B is essential for completion of the parasite life cycle.

Several hypotheses for the role of PbKIN8B can be envisaged.

1) The absence of PbKIN8B could cause defects at the level of the basal body, at the very beginning of the construction of the axoneme. Our IFA data show that PbKIN8B is present along the length of the axoneme, but we cannot exclude its presence already at the basal body. In *Plasmodium* and other apicomplexans such as *Toxoplasma*, the basal body appears as an amorphous electron dense structure in which only rarely a substructure composed of 9 singlet microtubules can be identified (Francia et al., 2015; Sinden et al., 1976). Due to the difficulty in resolving the substructure of the basal body it is impossible to comment on structural changes resulting from the absence of Pbkin8B. However, close observation of basal bodies and associated structures showed that fewer basal bodies can be observed in the mutant and those present have lost their tight association with nuclear poles (Table S3). Furthermore, it has been recently shown *in vitro* that, under certain conditions, B-microtubules can nucleate on A-microtubules, resulting in a doublet microtubules even in absence of a basal body (Schmidt-Cernohorska et al., 2019). Thus, even in the *ΔPbKIN8* mutant, singlet and doublet microtubules could potentially assemble without a functional basal body, as observed in the basal body mutant *Δsas-6* (Marques et al., 2015).

Interestingly, a default of cytoplasmic microtubules assembly, similar to the one observed in *ΔPbKIN8*, has been described after treatment of *P. berghei* gametocytes with azadirachtin, a plant limnoid and insecticide affecting male gametogenesis in *Plasmodium* (Billker et al., 2002). The authors suggest that azadirachtin would interact directly or indirectly with cytoplasmic components of the microtubule organizing center on the face of the spindle plaque, thereby disrupting both cytoplasmic microtubule patterning and the separation/rotation of the nuclear spindle poles at the prometaphase (Billker et al., 2002).

2) Assuming the basal bodies seen in the knockout mutant are functional and able to sustain the elongation of axonemal microtubules, the problem could lie in the stability of the assembled axoneme. As the axoneme grows, the abnormalities due to the absence of PbKIN8B cause the disruption of the connections between elements leading to a fatal dislocation of the structure. Several factors could exacerbate the instability:

First, coordinated activity of axonemal dyneins, located on the doublet microtubules causes sliding of microtubules during axonemal bending (for a recent review (Viswanadha, Sale, & Porter, 2017)). However, in *Plasmodium* fully formed axonemes start beating only seconds before exflagellation, limiting the sliding effect on the stability of the structure (Sinden & Croll, 1975).

Second, encasement of axonemes built on cellular protrusions enclosed in a membrane, as is the case in most eukaryotes, could stabilize the structure. This sheath is absent in *Plasmodium* due to its intracytoplasmic mode of assembly.

Third, *Plasmodium* operates with a reduced set of proteins for assembly and functioning of the axoneme. This structure, while fast to assemble in the cytoplasm, may be more susceptible to dislocation when just one component is missing.

On the other hand, PbKIN8B could play an indirect role and interact with various partners. Unfortunately, the small number of proteins involved in *Plasmodium* male gametogenesis identified and the absence of conservation of PbKIN8B in the most commonly used model organisms render the identification of putative partners difficult.

KIN8B could transport elements necessary for assembly or stabilization of the axoneme, that could be other motor proteins. In *Saccharomyces pombe*, the active kinesin-8 is a heterodimeric complex formed by KLP5 and KLP6 (Garcia, et al, 2002; Unsworth et al, 2008; West, et al, 2002). KIN8B in *P. berghei* displays several coiled-coil motifs adjacent to the motor domain, which could be involved in oligomerization.

PbKIN8B could also interact with one or several axonemal proteins among several motor proteins (as dyneins and kinesins, among which are kinesin-13 and −15, expressed in male gametogenesis) (Szklarczyk et al., 2015) (Invergo et al., 2017).

The phenotype observed in *ΔPbkin8B* resembles strongly the one of the basal body mutant *Δsas-6*, this protein – or another basal body protein – could therefore be a possible partner of PbKIN8B. This would be coherent with the IFA localization observed for *Pbkin8B-gfp*.

A concomitant study by Zeeshan et al., deposited on BioXriv, following the original submission of this paper, confirmed the localisation of PbKIN8B with the basal bodies and the axoneme, establishing a role of PbKIN8B in the basal body function (Zeeshan, Ferguson, et al., 2019). However, other roles in gametogenesis, which would explain the localisation of KIN8B, along the length of the axoneme, cannot be excluded. The lack of information on molecular actors involved in gametogenesis coupled to a complex and original system of axoneme assembly and gamete production, renders the attribution of partners to PbKIN8B complicated and future studies are necessary to dissect this crucial step in the parasite life cycle.

We present here the first characterization of kinesin 8B (PbKIN8B) involved in male gametogenesis in *P. berghei*. This study illustrates the importance of molecular motors in *Plasmodium* and shows that the absence of this kinesin causes severe defects that impede completion of the life cycle. Taken together, our results not only illustrate a previously unknown role for PbKIN8B in male gametogenesis, but also provide new insights into flagellar organization and function in *Plasmodium*.

## Experimental procedures

### Ethics statement

All animal work was carried out in accordance with the European regulations and in compliance with the French guidelines and regulations. The project was approved by the Ethic Committee CUVIER (authorization n°68-007).

### Generation of transgenic parasites

A pyrimethamine-sensitive clone of *P. berghei* NK65 strain (kindly provided by R. Ménard, Pasteur Institute, France) was used throughout this study to infect mice as described by Janse and collaborators (de Koning-Ward et al., 2000). *P. berghei* was maintained by cyclic passage in 4 to 6 weeks old female Swiss OF1 (Janvier labs, France). The *Pbkin8B-gfp* plasmid was generated by amplifying the final portion of the PbKIN8B coding sequence [nt1978-nt4378] with primers 111 and 112b (S1 Table). A unique restriction site for SacII in the middle of this region was used for single digestion and single crossover (vertical bar Fig S1A). The construct was inserted into the vector pl0016 (MRA-785 (BEI Resources)). The stop codon was removed and the gfp coding sequence was fused in-frame to the coding sequence. The plasmid also contained the *T. gondii* dhfr/ts resistance marker conveying resistance to pyrimethamine (Franke-Fayard et al., 2004). For PbKIN8B replacement (*ΔPbkin8B* lines), the transfection vector was sourced from the Sanger Institute (PbGEM-267699) (Schwach et al., 2015) and linearized by NotI prior to transfection. Details on tagging and knockout of PBANKA_020270 production are shown in Figs S1 and S2. Transfections were performed as described previously (Janse, Ramesar, & Waters, 2006). Following drug selection, two independent clonal populations of each genetic background resulting from two independent transfections were selected by limiting dilution and subsequent genotyping. Transfection experiments for gene disruption and gene tagging strategies were done in duplicate on different days using different batches of material.

### Genotypic analysis of mutants

The genotypes of *Pbkin8B-gfp* and *ΔPbkin8B* parasites were analysed by PCR with specific primers (Figs S1 and S2, Table S1). Briefly, for the C-terminal fusion *Pbkin8B-gfp* tagged parasites, integration was verified using primer 109 upstream of the amplified region and primer 115 in the *Pbkin8B-gfp* construct. Primers 53 and 54 served to confirm the presence of the resistance cassette. For the gene knockout parasites, two diagnostic PCR reactions were used as shown in Fig S2. Amplification by primers GT and GW2 was used to determine successful integration of the selectable marker at the targeted locus whereas primers QCR1 and QCR2 were used to verify deletion of the gene (S1 Table).

### Gametocyte preparation, exflagellation assays and ookinete conversion rates

Preparation of *P. berghei* gametocytes was realized as described previously (Beetsma, van de Wiel, Sauerwein, & Eling, 1998). Briefly, mice were injected intraperitoneally with 0.1 ml of 25 mg/l phenylhydrazine (to induce hyper-reticulocytosis) two days prior to infection by 10^7^ parasites. To reduce asexual parasitaemia, mice received sulfadiazine (10 mg/L) in their drinking water from day 5 to 7 after infection. On day seven, gametocyte-infected blood was collected for direct exflagellation assays or gametocyte purification before immunofluorescence assays, electron microscopy experiments or cytometry measurements as described by Billker *et al.* (Billker et al., 2004). After collection of whole blood on heparin, white blood cells were removed on CF11 cellulose (Whatman) columns. Gametocytes were separated from uninfected erythrocytes on a Nycodenz cushion made up from 48% Nycodenz stock (27.6% w/v Nycodenz in 5.0 mM Tris-HCl pH 7.2, 3.0 mM KCl, 0.3 mM EDTA) and RPMI1640 medium containing 25 mM HEPES (Sigma), 5% FCS, 4 mM sodium bicarbonate, pH 7.3. Gametocytes were harvested from the interface and washed three times in the appropriate buffer for the subsequent protocol. All manipulations were carried out at 19–22°C. Mature male and female gametocytes can be differentiated by gametocyte pigmentation following Giemsa staining (males appear blue and females pink).

For exflagellation assays, blood was diluted in 10 volumes of exflagellation medium (100 µM xanthurenic acid (XA) (Sigma) in RPMI 1640 (Thermo Fisher Scientific), pH 7.4). The actively moving gametes interacting with neighbouring RBC (exflagellation centres) were recorded 20 <u>m</u>inutes <u>p</u>ost <u>a</u>ctivation (mpa) by phase contrast microscopy in 10 x 1mm^2^ Malassez squares using a 40× objective. The number of exflagellation centres was then expressed relative to 100 male gametocytes for each sample. To perform the ookinete conversion assay, blood was first mixed at a 1:1 ratio in ookinete medium (RPMI pH8, 10% foetal calf serum, 100 µM XA). After 24 h to allow completion of gametogenesis and fertilization, ookinete conversion assays were performed as previously described (Tewari et al., 2005) by incubating samples with monoclonal antibody 13.1 (antibody against Pb28), conjugated with Cy3. The conversion rate corresponds to the proportion of ookinetes to all 13.1-positive cells (unfertilized macrogametes and ookinetes). Experiments were realized in biological triplicates.

### Western blotting

Asexual stages and gametocytes were isolated as described above. After the addition of Laemmli sample buffer, the samples were boiled and equal quantities of total protein were loaded on a 7 % SDS-polyacrylamide gel, before transfer to a nitrocellulose membrane (Thermo Fisher). Western blot analysis of *Pbkin8B-gfp* was performed under reducing conditions, using an anti-gfp rabbit antibody (1/3000, Abcam) coupled to an alkaline phosphatase conjugated goat anti-rabbit globulin (1/5000, Thermo Fisher).

### Mosquito infection

For mosquito infections, 3-to 8-day-old female adult *A. stephensi* mosquitoes were raised as previously described (Dimopoulos, Seeley, Wolf, & Kafatos, 1998). Day 3 post mouse-infection, mosquitoes were allowed to feed on anaesthetized infected mice for 20 min (Rodriguez et al., 2002; Sinden, Butcher, & Beetsma, 2002). Mosquitoes which had not fed, were discarded.

Engorged mosquitoes were dissected at day 8 or day 12 post feeding and the number of oocysts counted. For bite-back experiments, mosquitoes were infected with wild type or *ΔPbkin8B* parasites and after 21 days, 6-to 8-week-old female Tucks Ordinary mice were infected by the bite of these mosquitoes. After 5 days, parasitaemia was determined on mouse blood by blood smears and Giemsa staining and was followed for 14 days for mice infected by *ΔPbkin8B* parasites. All experiments were realized in duplicate. Differences between groups were calculated with Fisher’s exact test for prevalence and Mann-Whitney test for intensity.

### Indirect immunofluorescence assay (IFA)

Purified gametocytes were fixed at different time points in 3.7% (v/v) formaldehyde overnight at 4°C and processed as described previously (Becker et al., 2010). After a brief wash in PBS, cells were allowed to adhere onto poly-L-lysine coated slides and were then permeabilized by 0.5% (v/v) NP40 in PBS for 15 min. After a 30 minute saturation step, slides were incubated for 2 h with the first antisera (diluted 1/1000 for rabbit anti α–tubulin II, 1/2000 for rabbit anti-GFP, 1/200 for rabbit anti-spectrin, non-diluted for mouse TAT1 (Woods et al., 1989) followed by three washes before incubation for 1 h with appropriate fluorescently labelled secondary antibodies (Alexa Fluor®488 goat anti-mouse IgG (H+L), Alexa Fluor® 568 goat anti-rabbit IgG (H+L) both diluted 1/300 (Invitrogen)). After a 5 min DAPI incubation (5 microgram/ml), followed by a final wash, slides were mounted in Vectashield (Vector Laboratories). Parasites were visualized on a Nikon Eclipse TE 300 DV inverted microscope with a 100x oil objective mounted on a piezo electric device using appropriate fluorescence emission filters. Image acquisition (z-series) was performed with a back illuminated cooled detector (Photometrics CoolSnap HQ, 12 bit, RoperScientific, France) using a 0.20 µm step. Image processing was performed using Image J software (http://rsb.info.nih.gov/ij/).

### Electron microscopy

Gametocyte samples (described above) were fixed 15 and 30 mpa in 2.5% glutaraldehyde in 0.1 M phosphate buffer and processed for electron microscopy as previously described (Ferguson et al., 2005). Briefly, samples were post fixed in 1% osmium tetroxide, treated en bloc with 2 % uranyl acetate, dehydrated and embedded in epoxy resin. Thin sections were stained with uranyl acetate and lead citrate prior to examination in a JEOL1200EX electron microscope (Jeol UK Ltd).

### Flow cytometry

To measure the nuclear DNA content of activated gametocytes (wt, *ΔPbkin8B* and *Δcdpk4* -a mutant viable in blood infection that does not undergo DNA replication during microgametogenesis (Billker et al., 2004)) by flow cytometry, purified gametocytes were transferred into ookinete culture medium for activation of gamete formation. At 0, 8 and 15 mpa, cells were fixed overnight at 4°C in 0.04% glutaraldehyde. Blood of naïve mice was fixed in the same conditions and used as a negative control. DNA was stained with YOYO-1 (Bouillon, Gorgette, Mercereau-Puijalon, & Barale, 2013). Briefly, gametocytes and blood samples were treated for 10 minutes with 0.25% TritonX100 in PBS after centrifugation at 100g for 5 minutes. They were then treated with 0.05 mg/ml RNase A/T1 (Thermo Scientific(tm)) for 4 h at 37°C. Finally, DNA was stained with 10.24 µM YOYO-1 (Invitrogen(tm)). A gametocyte sample was duplicated for an unstained control. After overnight incubation at 4°C, the supernatant was replaced by FACSFlow solution and flow cytometry data acquired on a FACSVerse (BD Biosciences). Data processing and analysis were performed using the FACSuite Software v 1.0.5 (BD Biosciences). YOYO-1 fluorescence was excited with a blue laser (20 mW) at 488 nm and the signal was detected at 527 +/-16 nm. Quality control based on FSC-A vs SSC-A and FSC-H vs FSC-A gating was applied to remove debris and doublets respectively. Then, histograms of YOYO-1-A signal (in log scale) were analysed to determine nucleic acid content. Gates were established resulting in populations P1 corresponding to the complete YOYO+ population, P2 composed of cells with the lowest DNA content, and P3, corresponding to cells with the highest DNA content. For each sample the FACS profile was established and the ratio of populations (P2 and P3) was expressed as a percentage of P1.

## Acknowledgements

We wish to thank Joy Alonso and Nathalie Dogna for animal care, Elisabeth Mouray, Lisy Raveendran and Cyril Willig (CeMIM Platform of the MNHN) for technical help and Philippe Bastin (Institut Pasteur, Paris) for critical reading of the manuscript and helpful discussions.

This project was funded by ATM from Muséum National d’Histoire Naturelle, Paris.

## Supplementary material

**Table S1.**
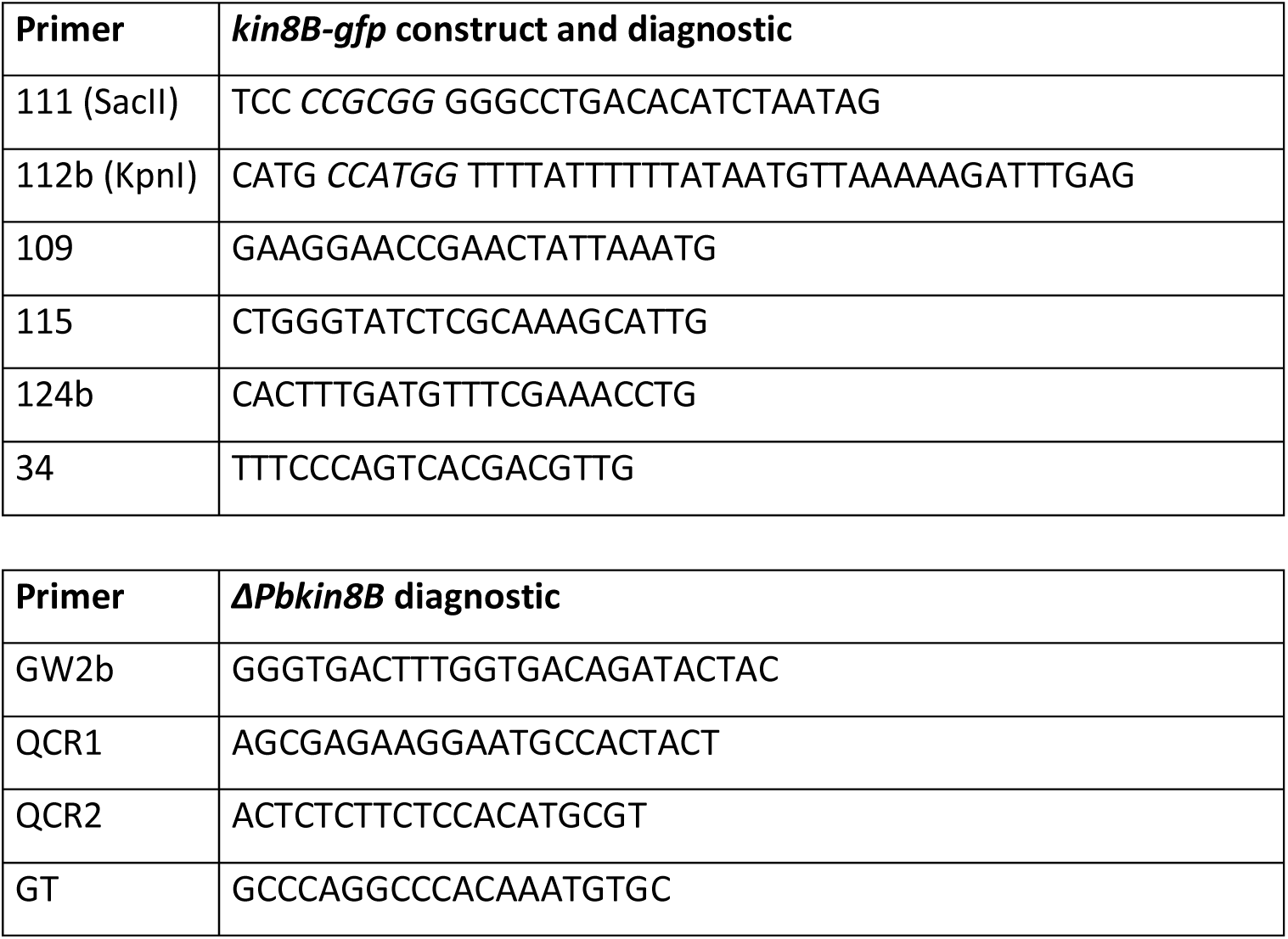
Primer sequences for construction of *Pbkin8B-gfp* vector and verification of integration of *Pbkin8B-gfp*, respectively *ΔPbkin8B*, in the *P. berghei* genome.

**Table S2.**
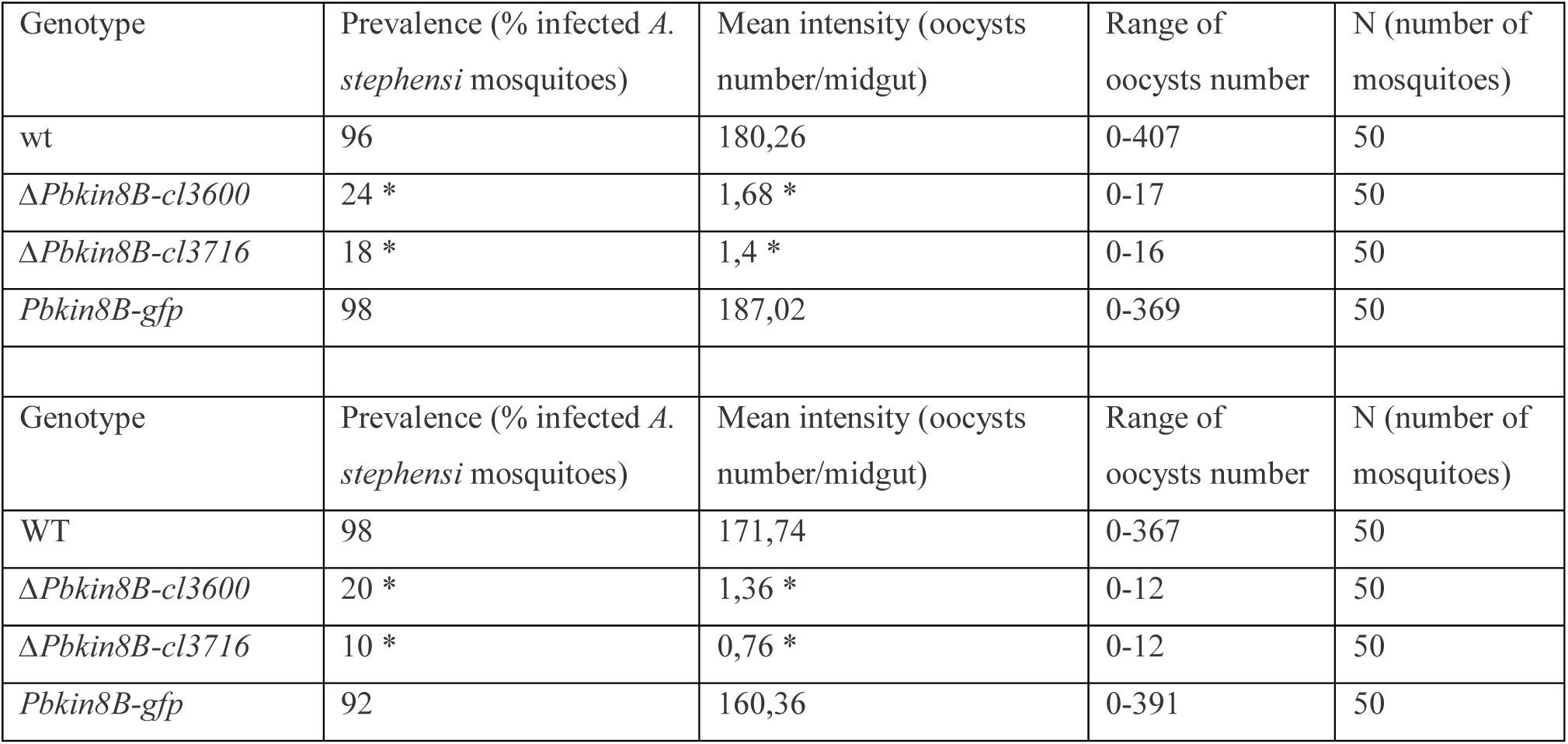
Prevalence and Intensity of mosquito infections. For each replicate, mice with similar parasitaemias were fed to mosquitoes. Fully fed mosquitoes were kept until midgut dissection, midguts were mounted on slides and oocysts were counted under the microscope. Differences between groups were calculated with Fisher’s exact test for prevalence, Mann-Whitney test for intensity. Asterisk Ȫ indicate statistically significant differences with p-values lower than 0.0001 and 0.005, respectively.

**Table S3.**
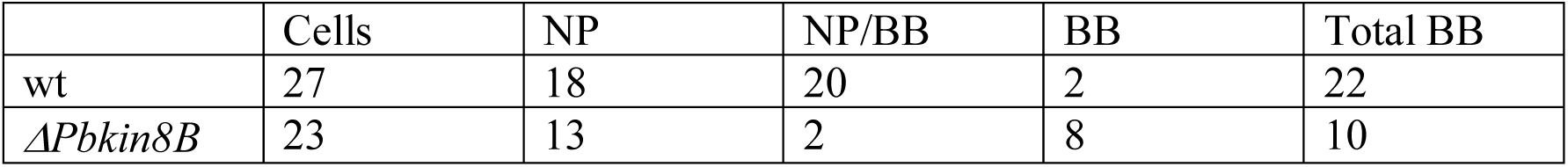
Number of nuclear poles, number nuclear pole with basal bodies and number of basal bodies without nuclear poles. Quantification of microtubule organising centres (nuclear poles and basal bodies) in wt and ΔPbkin8B gametocytes were counted in 27 wt and 23 ΔPbkin8B and gametocytes.

**Fig S1.**
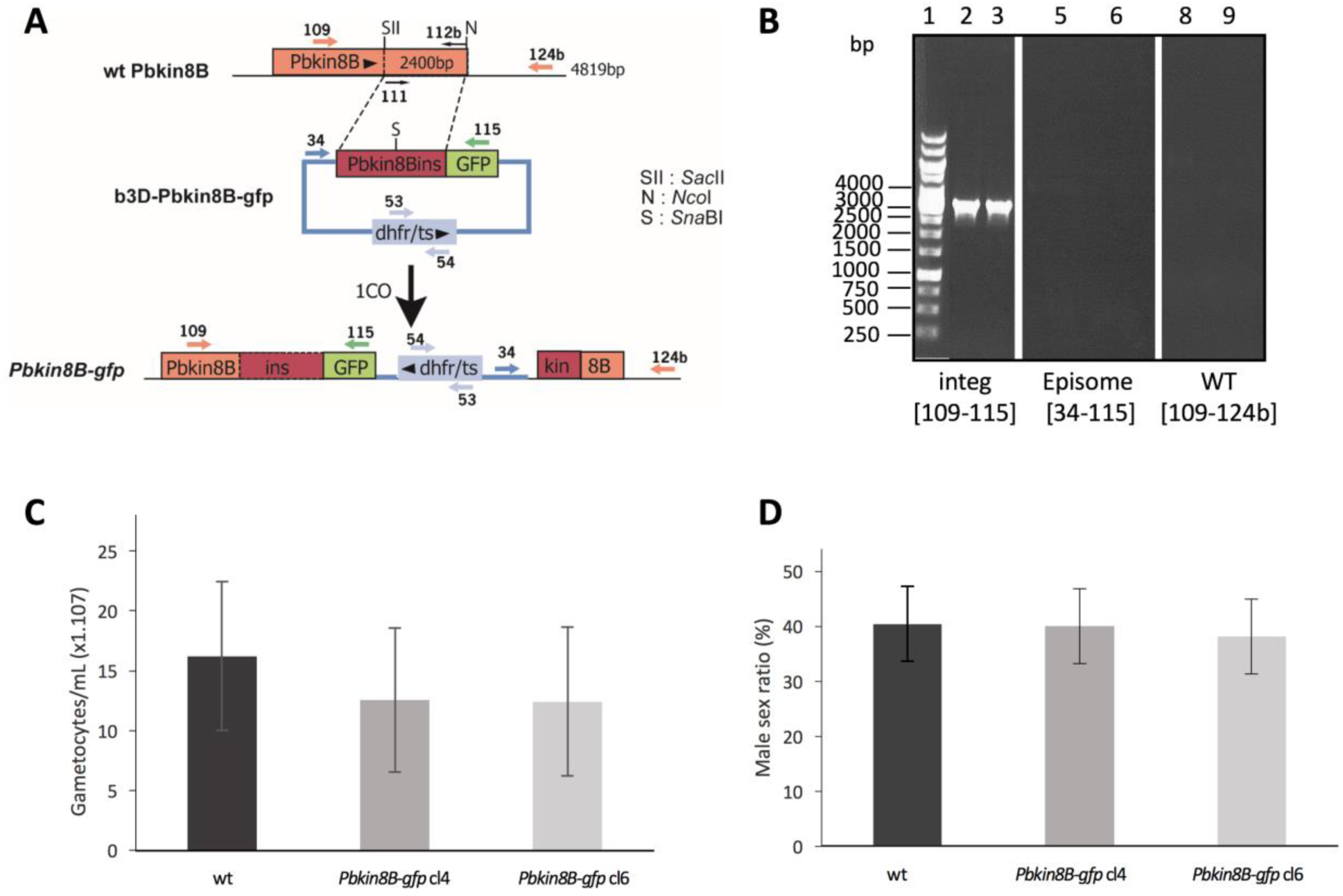
Generation and genotypic analysis of *Pbkin8B-gfp* parasites. (A, B) Generation and genotypic analysis of Pbkin8B-gfp parasites. (A) Schematic representation of Pbkin8B locus before and after insertion of a plasmid containing a partial Pbkin8B sequence fused to a gfp tag. Single homologous recombination occurred between the plasmid and the homologous region of the 3’-terminal part of the genomic locus of Pbkin8B. Selection was realized using the Toxoplasma gondii dhfr/ts resistance marker present in the plasmid. (B) Analysis of genomic DNA from Pbkin8B-gfp (clones 4 and 6) and wt parasites by PCR. PCR amplifications were realised to verify integration of the construct in the correct locus using primers detailed in Supplemental Table 1. Lanes 2, 5 and 8 correspond to clone 4; lanes 3, 6 and 9 correspond to clone 6. PCR amplification confirmed correct integration of the construct (lanes 2 and 3), absence of wt genotype (lanes 5 and 6) and of episomal plasmid (lanes 8 and 9). (C, D) Phenotypic analysis of gametocyte production of Pbkin8B-gfp and wt on Giemsa stained blood smears. (C) The number of gametocytes was determined for 4 infected mice per genotype. (D) Male sex ratio of Pbkin8B-gfp clones 4 and 6 is similar to wt. Differences between groups were not statistically different. SD are reported as bars on the figures.

**Fig S2.**
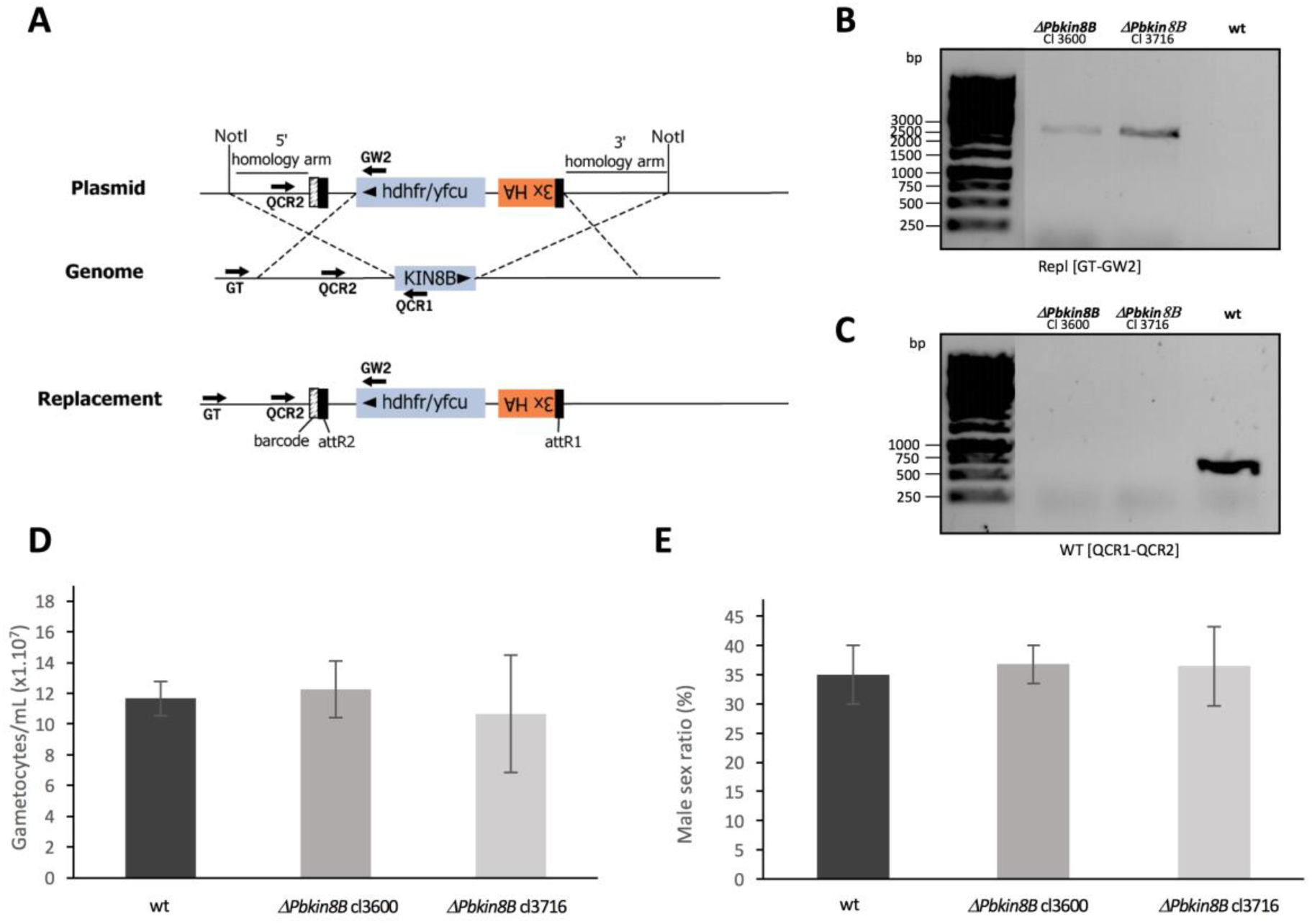
Genotypic and phenotypic analysis of *ΔPbkin8B* parasites. (A-C) Generation and genotypic analysis of ΔPbkin8B parasites. (A) Schematic representation of Pbkin8B locus before and after replacement of the coding sequence by human dihydrofolate reductase thymidilate synthase / yeast cytosine deaminase and uridyl phosphoribosyl transferase (hdhfr/yfcu) gene which confers resistance to pyrimethamine through homologous recombination with the 5′ and 3’ homology arms of Pbkin8B (PBANKA_020270). Primer positions for verification of replacement are indicated by arrows. The transfection vector sourced from the Sanger Institute (PbGEM-267699). (B, C) Analysis of genomic DNA from ΔPbkin8B (clones 3600 and 3716) and wt parasites by PCR. (B) PCR amplifications were realised to verify integration of the construct in the correct locus using primer GT and primer GW2. (C) Amplifications with primers QCR2 and QCR1 confirm the absence of Pbkin8B gene in the two ΔPbkin8B clones. (D, E) Phenotypic analysis of gametocyte production of ΔPbkin8B and wt on Giemsa stained blood smears. (D) The number of gametocytes was determined for 4 infected mice per genotype (i.e wt, ΔPbkin8B clone 3600 and clone 3716). (E) Male sex ratio of ΔPbkin8B (clones 3600 and 3716) is similar to wt. Differences between groups were not statistically different. SD are reported as bars on the figures.

**Fig S3.**
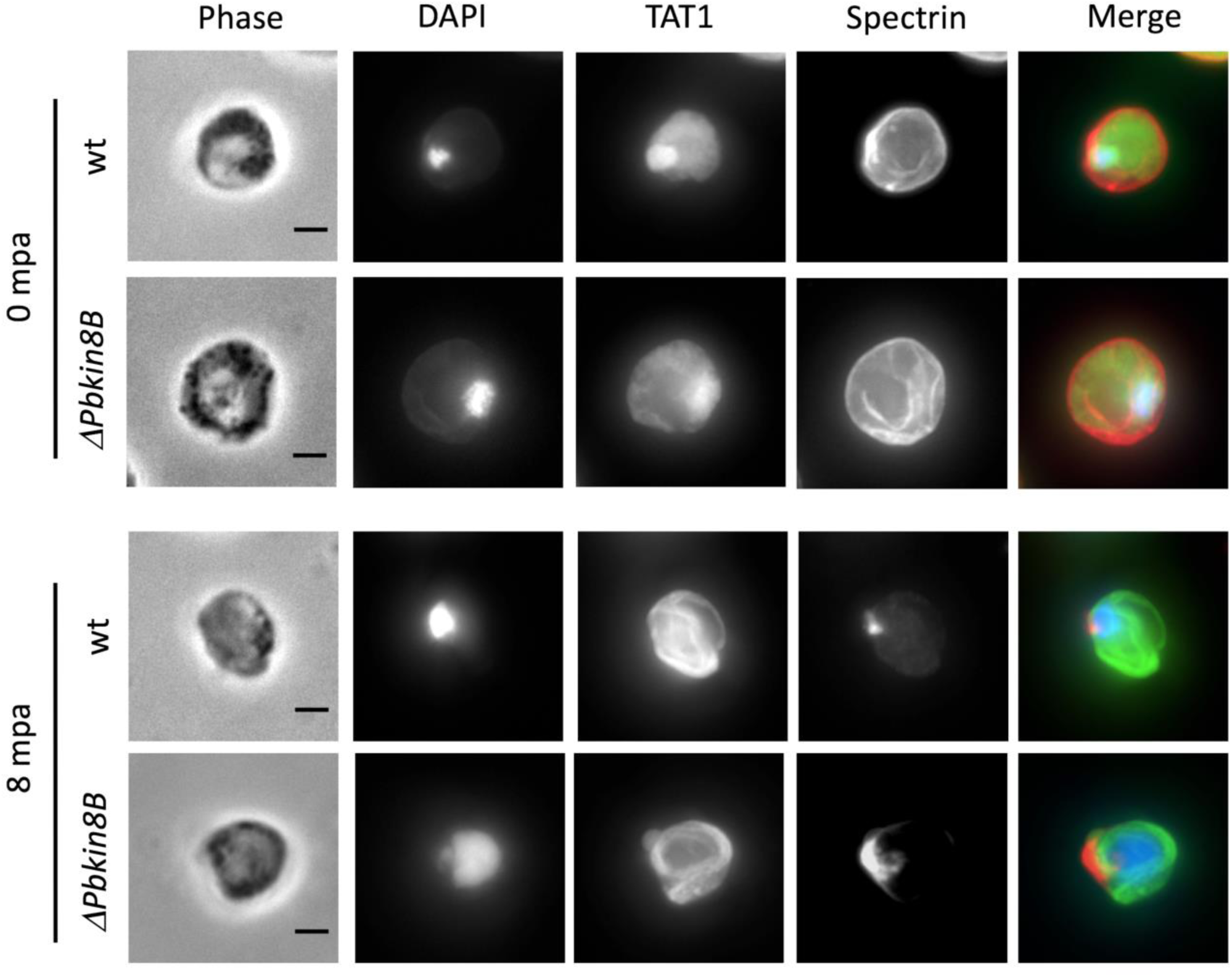
After activation *ΔPbKIN8B gametocytes* egress quickly from red blood cells. Immunofluorescence assay of wt and ΔPbkin8B male gametocytes at 0 and 8 mpa. DAPI staining of DNA is seen in blue, TAT1 in green and anti-spectrin in red. Immunofluorescence images correspond to the maximum intensity projection of the z-series. At 8 mpa, the erythrocyte membrane is degraded compared to 0 mpa, while microtubules have formed in the cytoplasm. Scale bar: 2 µm.

